# Machine learning prediction of particle-size distribution from infrared spectra, methodologies and soil features

**DOI:** 10.1101/2020.05.04.076471

**Authors:** Elizabeth J. Parent, Serge-É. Parent, Léon E. Parent

## Abstract

Accuracy of infrared (IR) models to measure soil particle-size distribution (PSD) depends on soil preparation, methodology (sedimentation, laser), settling times and relevant soil features. Compositional soil data may require log ratio (*ilr*) transformation to avoid numerical biases. Machine learning can relate numerous independent variables that may impact on NIR spectra to assess particle-size distribution. Our objective was to reach high IRS prediction accuracy across a large range of PSD methods and soil properties. A total of 1298 soil samples from eastern Canada were IR-scanned. Spectra were processed by Stochastic Gradient Boosting (SGB) to predict sand, silt, clay and carbon. Slope and intercept of the log-log relationships between settling time and suspension density function (SDF) (R^2^ = 0.84-0.92) performed similarly to NIR spectra using either *ilr*-transformed (R^2^ = 0.81-0.93) or raw percentages (R^2^ = 0.76-0.94). Settling times of 0.67-min and 2-h were the most accurate for NIR predictions (R^2^ = 0.49-0.79). The NIR prediction of sand sieving method (R^2^ = 0.66) was more accurate than Bouyoucos (R^2^ = 0.53). The NIR 2X gain was less accurate (R^2^ = 0.69-0.92) than 4X (R^2^ = 0.87-0.95). The MIR (R^2^ = 0.45-0.80) performed better than NIR (R^2^ = 0.40-0.71) spectra. Adding soil carbon, reconstituted bulk density, pH, red-green-blue color, oxalate and Mehlich3 extracts returned R^2^ value of 0.86-0.91 for texture prediction. In addition to slope and intercept of the SDF, 4X gain, method and pre-treatment classes, soil carbon and color appeared to be promising features for routine SGB-processed NIR particle-size analysis.

*Soil Classification* (*Soil Taxonomy*): Inceptisols, Spodosols

## 1. Introduction

Soil particle-size distribution is of prime importance for plant growth and soil management [1]. Mimicking particle sedimentation in natural water bodies, PSD has been traditionally quantified using the sieve-pipette method that determines particle mass, and the sieve-hydrometer or sieve-plummet balance method that measures changes in suspension density [2]. Sedimentation techniques are thought to overestimate the concentration of plate-like clay particles that do not fit into Stokes’ law. Reynolds’ number should be less than 0.05, otherwise, the drag force for sedimentation increases faster than predicted by Stokes’ law [3]. Because organic matter binds soil particles [4], soils may be pre-treated with peroxide to destroy organic matter and disperse soil particles. Laser techniques are much faster than sedimentation techniques but tend to underestimate clay-size particles due to insufficient particle dispersion despite sonication pre-treatment, and this may require using conversion equations [5,6].

On the other hand, visible and near infrared (VIS–NIR) spectroscopy is an efficient soil quality and fertility screening tool [7] because spectra correlate well with several chemical, physical and mineralogical properties [8]. The VIS represents the visible light range between 350 and 780 nm, from violet to red. Because most routine soil and plant laboratories are equipped with NIR spectrometers for forage analysis, they may contribute to documenting soil characteristics at low cost. Where VIS is not available, soil RGB (red-green-blue) can be assessed from the Munsell color chart using computer models. Mid-infrared (MIR) is generally more accurate than NIR [9] but requires more sample preparation, limiting its application as routine determination method.

Infrared spectroscopy (IRS) more accurately predicts clay than sand and silt contents [10] because the IR spectrum is sensitive to clay mineralogy [10–12] and total reflectance decreases as grain size increases [13,14]. Light absorption is also influenced by soil features such as particle roundness [15–18], soil pH, and the Mehlich-3 soil test for Ca, Mg and Mn [19]. The IRS detects Al-OH (2200 nm) and Fe-OH (2290 nm) [20,21] that in turn have an impact on soil structure [22] and IR reflectance [23]. The VIS-NIR spectra are sensitive to soil moisture and C content [24]. Organic matter, multi-nutrient extraction and reconstituted bulk density from scooped soil samples are also common features quantified in routine laboratories.

The sedimentation methods provide percentages of sand, silt, and clay from the log-log relationship between settling time and suspension density. The slope and intercept return proportions of sand, silt and clay at pre-selected settling times that may vary between laboratories [1], thus affecting the accuracy of IRS models. Providing more flexibility using the slope and intercept of the log-log relationship and reducing the arbitrariness of the settling time selection may allow increased reliability of IRS models calibrated against Bouyoucos methods.

There are unattended sources of error in IRS calibration. There is systematic negative covariance between sand, silt and clay fractions due to resonance within the ternary diagram [25]. Indeed, there are D-1 degrees of freedom in a D-part composition [26]. Not considering the problem of closure to 100% in statistical analysis, confidence intervals about means of proportions may take values outside of the compositional space, i.e. < 0 or > 100% [27], and the measures of distance and dissimilarity are non-Euclidian [28]. To return unbiased statistical results, orthonormal balances among subsets of components can be computed as D-1 isometric log ratios (*ilr*) [29]. The back-transformed *ilr* values allow recovering proportions of sand, silt, and clay totalling exactly 100% within the limits of the ternary diagram.

For both the sedimentation and laser techniques there are several soil pre-treatments (peroxide, sodium hypochlorite, sodium hexametaphosphate, sonication intensity), soil features, calibration techniques, options (NIR 2X, NIR 4X, settling times or suspension density function for sedimentation; pump and stirrer spin, refractive index of the medium, real or imaginary refractive index, density for laser) and expressions (percentages, ratios) that influence results of particle-size distribution. Machine learning (ML) is an emerging data mining technique of artificial intelligence that can unravel patterns and rules in large data sets [30] and predict target variable from input data [31]. Machine learning methods can account for numerous independent variables that may impact on NIR spectra to assess accurately soil particle-size distribution.

This paper is presented in two parts, one focusing on methodology and the other on modeling. In the methodology section, we hypothesized that methods and pre-treatments return different results and thus cannot be combined to run a single IRS calibration set without input information on methodologies. In the modeling section, we hypothesized that: 1. IRS cannot accurately predict PSD from laser methods due to underestimation of the clay fraction; 2. IRS accurately predicts sand fraction determined using sand sieving methods; 3. IRS more accurately predicts PSD from sedimentation methods after accounting for soil features; 4. IRS prediction accuracy for PSD is improved using the slope and intercept of the log-log relationship and isometric log ratios compared to raw percentages and pre-determined settling times; and 5. MIR prediction accuracy for PSD is higher compared to the NIR spectra. The objective of this study was to reach high IR-ML model accuracy for soil texture determination integrating numerous soil features, determination methods and methodological modifiers.

## 2 Materials and methods

### 2.1. Characterization of soils

The data set of 1298 soil samples collected in the arable layer (0-20 cm) was obtained from several research institutions in Québec, Canada. The main crops were maize (*Zea mays*) cereals and forages on coarse-to fine-textured soils, potatoes (*Solanum tuberosum*) on sandy loams and loamy sands, and cranberries (*Vaccinium macrocarpon*) on sandy soils. The soils were mainly Inceptisols and Spodosols.

Soil samples were air-dried then passed through a 2-mm sieve [32,33]. The 2-mm sieved soil was 3-mL scooped and then weighed to determine the reconstituted bulk density as performed routinely in soil testing laboratories. Soil color was assessed on dry samples using the Munsell chart and then transformed into RGB percentages using “munsell2rgb” in R. The total carbon (Ct) was quantified using the Leco CNS analyzer (Leco Corporation, St. Joseph, Michigan). For pH determination, 10 g of soil was mixed with 20 mL 0.01M CaCl_2_. For oxalate extracts, a 0.5 g sample of soil was mixed with 20 mL of oxalate solution (0.2 M of ammonium oxalate and 0.2 M of oxalate acid) and agitated for 4 h in the dark (34). The mixture was centrifuged at 2000 rpm for 5 min and then filtered through Whatman no. 40 paper. Concentrations of P, Fe, Ca, Al, Mn and Si were quantified by ICP-OES. Soils were also extracted using the routine Mehlich3 method [35]. Concentrations of P, Ca, Mg, Fe, Al, Mn, Zn and Cu were quantified by ICP-OES.

### 2.2 Particle-size analysis

Particle size distribution was analyzed in 50-100 g samples using the hydrometer method [36]. A separate batch of soils was pre-treated with peroxide for comparison with the no-peroxide pre-treatment. Samples were mixed with 0.05 M hexametaphosphate and agitated at 300 rpm for 16 h. The mixture was transferred to a 1 L cylinder and hand-shaken for 30 sec. Suspension density readings (g L^-1^) taken after 0.75-, 5-, 120-, 420-, and 1440-min were referred to as the Bouyoucos multi-point method. Samples reporting suspension density after 0.67 and 120 min as originally suggested [37] were assigned to the 2-point 2-h Bouyoucos method. The clay fraction was also recorded after 7-h settling time, close to the 6-h settling time used by Gee and Bauder (1979). Although the clay fraction was overestimated as compared to the 7-h Bouyoucos multi-point method, the 2-point 2-h Bouyoucos method was selected as our reference because it has been widely used as a proximate method in soil surveys combined with tactile assessment. After taking the last reading, the entire hydrometer contents were passed through 1-, 0.5-, 0.25-, 0.10- and 0.05-mm sieves under tap water to clean the coarser particles of any adhering finer particles and to determine the sand-size distribution. The suspension density was transformed into particle-size percentages by mass using standard equations [37,38]. The Bouyoucos curve relating suspension density to settling time was log-transformed (ln) to determine the slope and intercept as model parameters.

Samples were also analyzed using the Mastersizer 2000 Laser particle size analyzer (Malvern Instruments, Worcestershire, UK, measurements at 633 nm and 466 nm) combined with Hydro 2000G (800-mL tap water volume, 500-rpm stirrer and 2000-rpm pump) with or without ultrasonic action at nominal 40 kHz frequency for 2 min [39]. Only samples within 10 to 20 % of obscuration were retained in the database. The refractive index of the medium was set at 1.5.

### 2.3 Calculation

The three particle-size fractions and carbon content were isometric log ratio (*ilr*) transformed as follows [29]:

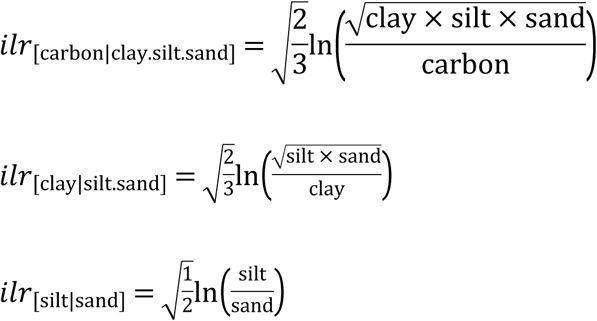

### 2.4 Spectral data acquisition

The air-dried and 2-mm sieved samples [40] were placed into a 5-cm quartz cup then scanned using a Nicolet Antaris FT-NIR analyzer (Thermo Electron Corp., Ann Arbor, Michigan). Absorbance was measured with gains of 2X or 4X. Triplicated spectra were scanned 30 times in the range of 9090 to 4000 cm^-1^ (1100 to 2500 nm) at a resolution of 2 cm^-1^ (0.3 nm at 1250 nm) [41]. A library of 3069 NIR spectra with 2X gain and 668 spectra with 4X gain were obtained after probabilistic quotient normalization and first derivation [42]. All samples were stored in white plastic storage containers.

The soil and KBr samples were ground to less than 74 µm using a mortar and pestle [43], then oven-dried at 105°C for 3 h [44]. A 2 g sample was weighed and mixed with 200 g KBr powder. The mixture was pressed at 517 MPa for 3 min using a manual hydraulic press (Carver, Model 4350.L, Carver Inc., Wabash, Indiana) to yield clear 13-mm-diameter pellets. The mid-infrared spectra were scanned using a transmission DTGS detector Varian 1000 FT-IR Scimitar series spectrometer (Varian Inc., Palo Alto, California). Spectra were in the range of 4000 to 400 cm^-1^ (2500 to 25 000 nm) at 4 cm^-1^ (0.6 nm) resolution. Each pellet was scanned 10 times and rotated once manually at 90°. A library of 413 MIR spectra was obtained.

### 2.5 Spectral data modeling

Statistical analyses were conducted using the R-3.6.1 version [45] with the packages “tidyverse”, “reshape2”, “stringi”, “signal”, “mvoutlier”, “caret”, “compositions”, “soiltexture”, “aqp”, “patchwork”, “broom”, “ggExtra”, and “outliers”. The machine learning method was GBM (Stochastic Gradient Boosting). The NIR and MIR spectra were normalized and passed through a binning process with 10 cut points and a Savitsky-Golay smoothing filter. After first derivation, spectra compressed into scores using principle component analysis (PCA) [46]. Because of the diversity of the soil samples, we did not remove outliers. Sand, silt and clay percentages, oxalate and Mehlich-3 extracts were *ilr*-transformed. Every set was divided into training (70%) and testing (30%). The R code elaborated by Elizabeth Parent and Serge-Etienne Parent is available at: https://github.com/eliparent/Texture.

## 3 Results

### 3.1 Soil characterization

There was a large spectrum of soil properties (Table 1). In comparison, soils of the region have been reported to range from 5 to 137 g organic matter kg^-1^, 560 to 19 992 mg Fe_oxalate_ kg^-1^ and 270 to 36 638 mg Al_oxalate_ kg^-1^, and 4-494 g sand kg^-1^, 40-734 g silt kg^-1^, and 6-796 g clay kg^-1^ [47]. Particle-size distribution is presented in Fig. 1. Orthonormal balances were normally distributed except for [clay | silt,sand] using the laser method (Fig. 2). Indeed, the clay content was systematically underestimated, and the silt content systematically overestimated by the laser method.

**Table 1.** Ranges of soil properties (0-20 cm) in the data set (particle-size distribution according to the multi-point 7-h Bouyoucos method using 45-sec settling time for sand and 7-h settling time for clay)

| Site | Mean | Standard deviation | Minimum | Maximum |
| --- | --- | --- | --- | --- |
| pH (0.01 M CaCl<sub>2</sub>) | 5.36 | 0.67 | 3.37 | 7.90 |
|  |  | g kg<sup>-1</sup> |  |  |
| Sand total | 613 | 24 | 1 | 986 |
| • 1-2 mm | 41 | 50 | 0 | 282 |
| • 0.5-1.0 mm | 108 | 115 | 1 | 541 |
| • 0.25-0.5 mm | 191 | 126 | 5 | 580 |
| • 0.1-0.25 mm | 235 | 154 | 3 | 766 |
| • 0.25-0.01 mm | 118 | 103 | 0 | 587 |
| Silt | 256 | 15 | 5 | 816 |
| Clay | 131 | 13 | 5 | 839 |
| Total C | 25.5 | 2.4 | 0.1 | 443 |
| Total N | 1.9 | 1.6 | 0.7 | 19 |
| Total S | 0.3 | 0.4 | 0 | 10.3 |
|  |  | mg kg<sup>-1</sup> |  |  |
| Fe<sub>oxalate</sub> | 5599 | 2449 | 447 | 10400 |
| Al<sub>oxalate</sub> | 4000 | 2693 | 528 | 15346 |
| Mn<sub>oxalate</sub> | 309 | 304 | 7 | 2287 |
| Ca<sub>oxalate</sub> | 30 | 14 | 3 | 153 |
| P<sub>oxalate</sub> | 803 | 464 | 68 | 2266 |
| Si<sub>oxalate</sub> | 829 | 578 | 88 | 5585 |
| P<sub>Mehlich3</sub> | 118 | 62 | 6 | 362 |
| Ca<sub>Mehlich3</sub> | 1776 | 1437 | 113 | 5103 |
| Mg<sub>Mehlich3</sub> | 191 | 224 | 5 | 1572 |
| Fe<sub>Mehlich3</sub> | 232 | 108 | 72 | 715 |
| Al<sub>Mehlich3</sub> | 1265 | 431 | 310 | 2241 |
| Mn<sub>Mehlich3</sub> | 25 | 20 | 5 | 206 |
| Zn<sub>Mehlich3</sub> | 8 | 9 | 0 | 130 |
| Cu<sub>Mehlich3</sub> | 4 | 3 | 0.4 | 30 |
|  |  | % |  |  |
| Green | 47 | 8 | 28 | 68 |
| Blue | 34 | 10 | 11 | 58 |
| Red | 54 | 7 | 31 | 75 |
|  |  | g cm<sup>-3</sup> |  |  |
| Reconstituted bulk density | 1.06 | 0.16 | 0.71 | 1.46 |

**Fig. 1.**
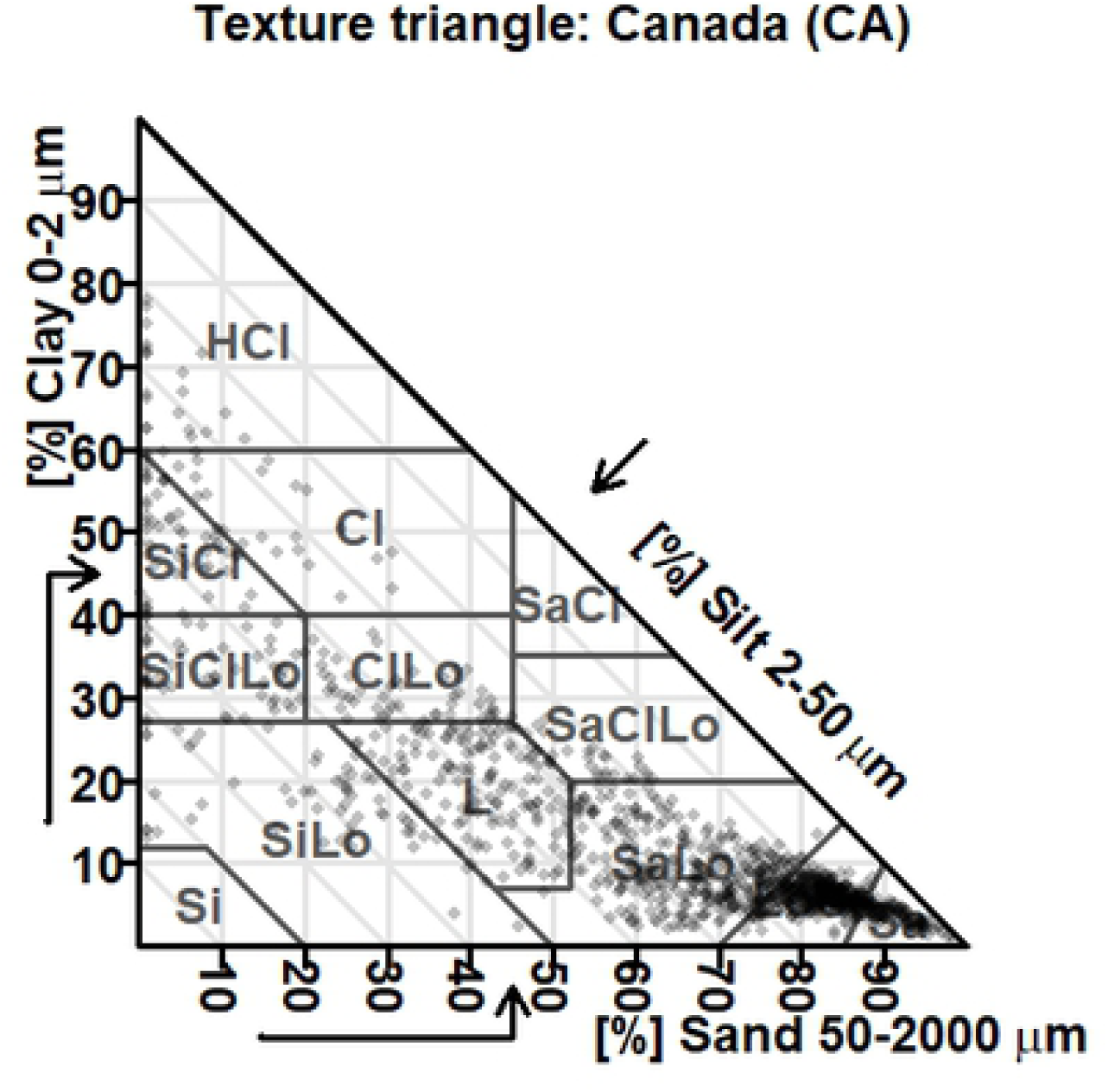
Particle size distribution of studied soils in the Canadian textural diagram.

**Fig. 2.**
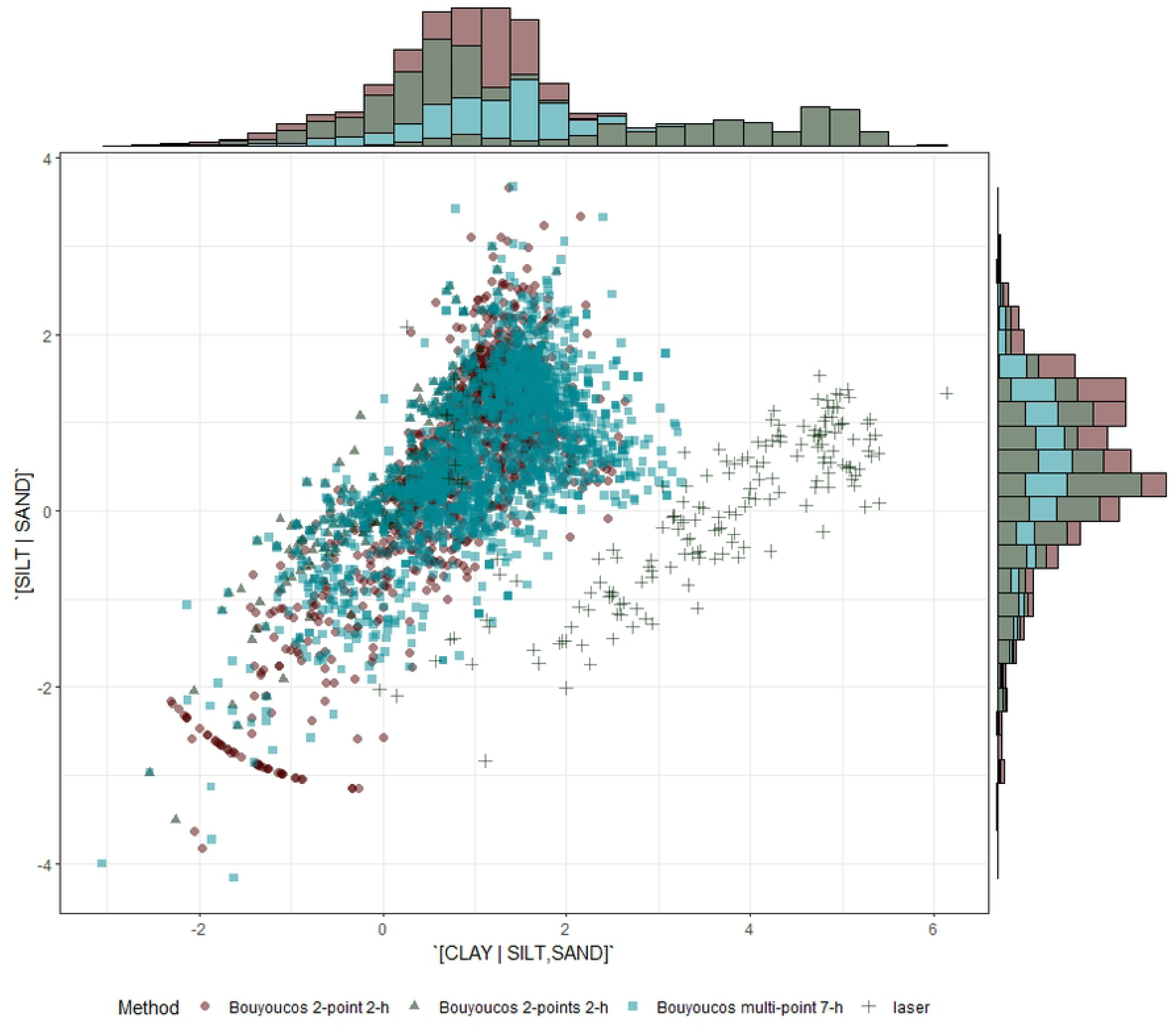
Distribution of ilr textural variables across methodologies.

### 3.2 Methodologies

Paired t-tests comparing methods are presented in Table 2. Comparisons involving the peroxide pre-treatment must be interpreted with care due to the smaller number of observations (26 to 38). Mean *ilr* differences between the reference 2-point 2-h Bouyoucos method and other methods are illustrated in Fig. 3. The difference between the laser and the 2-point 2-h Bouyoucos methods depended on the *ilr* coordinate (also shown in Fig. 2). After 2-min of sonication, the [silt | sand] balance differed slightly between methods while the [clay | silt,sand] balance differed markedly between methods, indicating insufficient dispersion of clay-size particles.

**Table 2.**
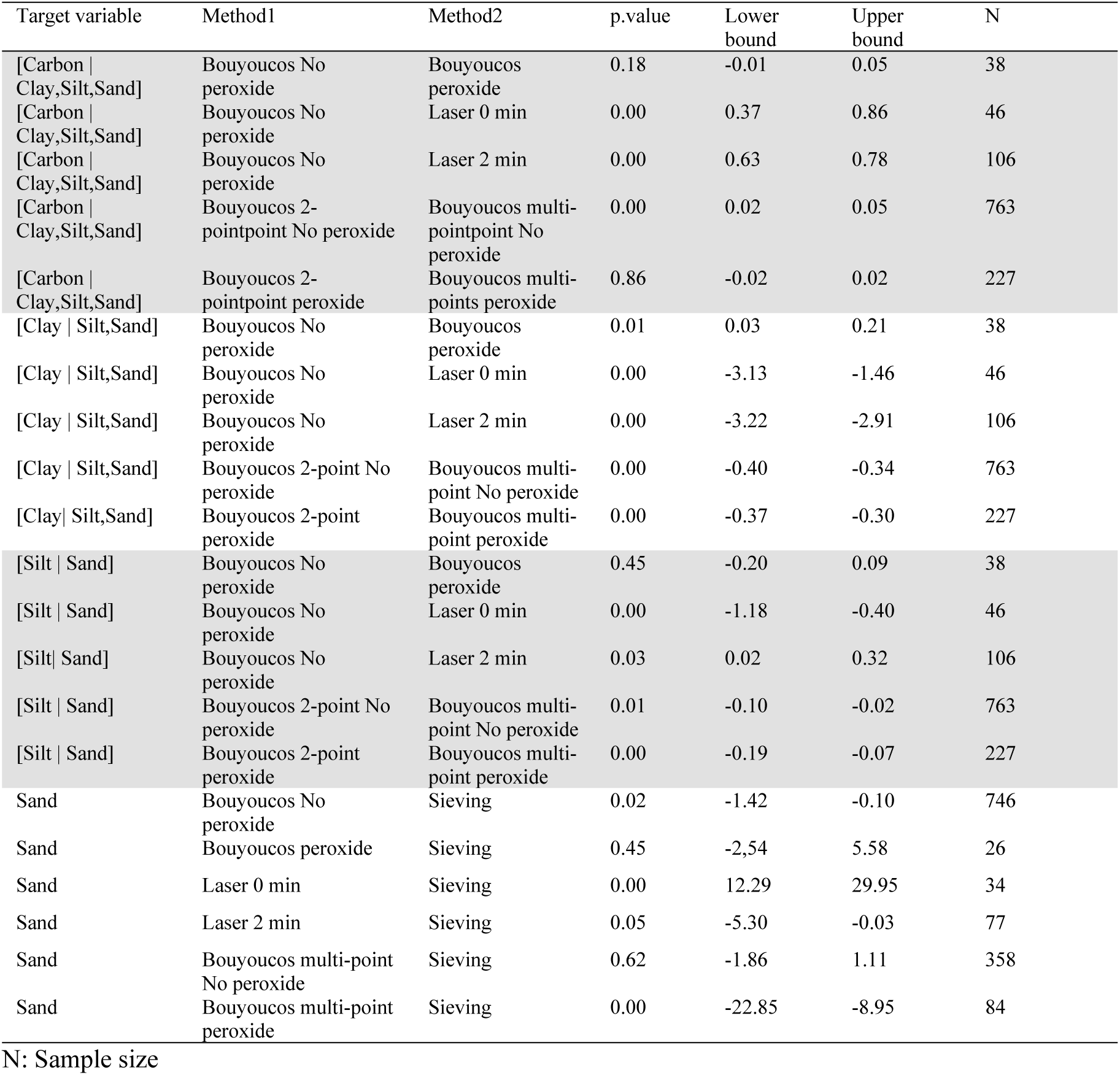
Comparison of methods (method1 minus method2) using paired t-test and confidence intervals (p ≤ 0.05)

**Fig. 3.**
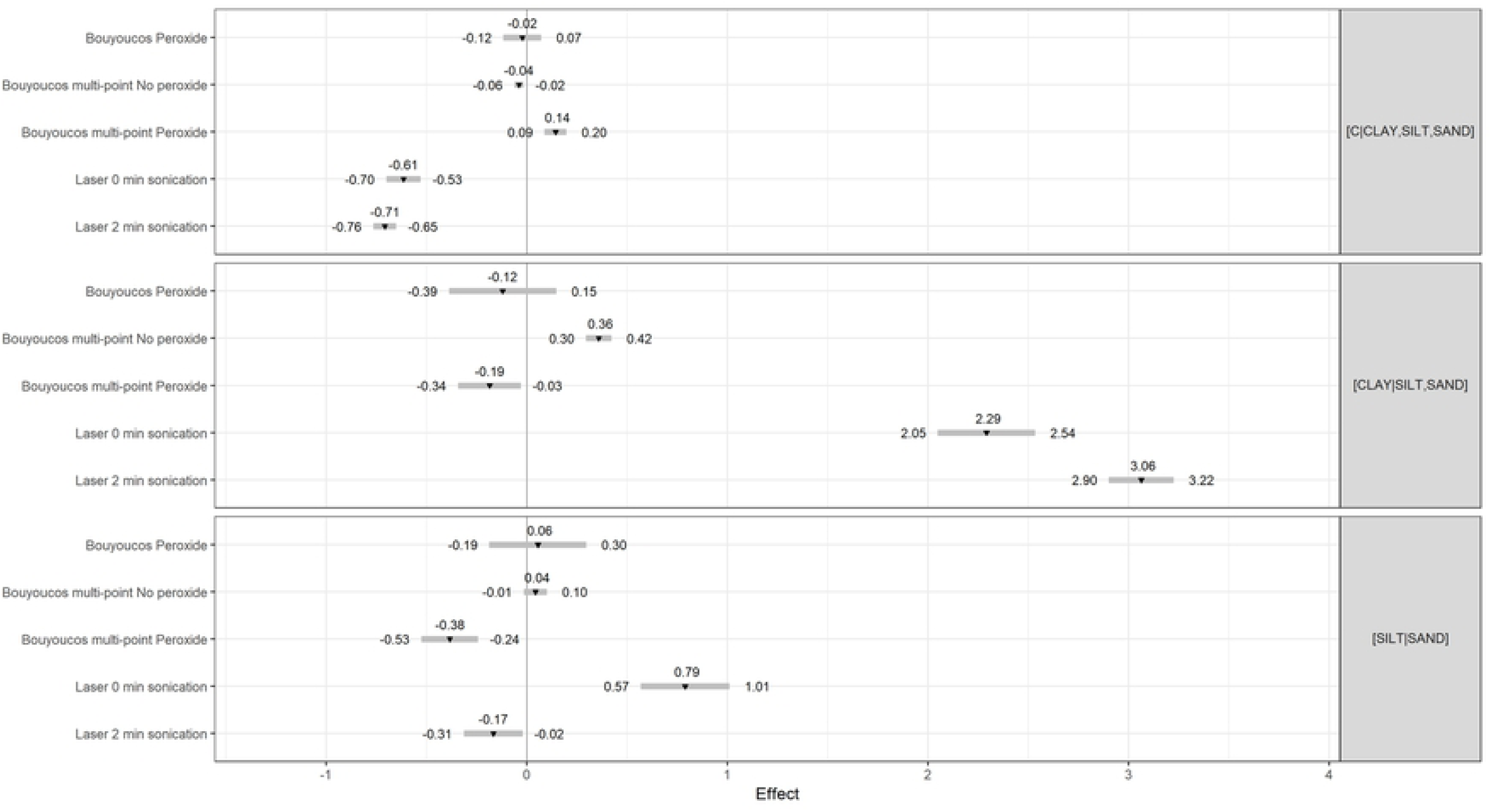
Mean differences (p ≤ 0.025) in textural balances between several methodologies against the reference 2-point 2-h Bouyoucos method without pre-treatment.

As expected, the multi-point 7-h Bouyoucos method returned a smaller proportion of clay-size particles compared to the 2-point 2-h Bouyoucos method due to longer settling time. The peroxide pre-treatment for the multi-point 7-h Bouyoucos method tended to increase clay and silt fractions. The [silt | sand] balance resulting from the no-peroxide pre-treatment preceding sedimentation methods was comparable across methods, with the exceptions of the laser and pretreated multi-point 7-h Bouyoucos methods. Excluding the 0-min sonicated laser and comparable methods, the amount of sand was slightly lower compared to sand sieving. Sand contents did not differ significantly between the peroxide pre-treatments or multi-points calculation without pre-treatment. Except for the peroxided pre-treated dataset, the [carbon | clay,silt,sand] balances were differentially influenced by methodologies.

### 3.3 Spectral data modeling

To conduct predictions to details of methodologies and features, the data set was divided into eight subsets (Table 3). Prediction models were run for each subset. In general, prediction accuracies were weakest for the silt fraction and were similar whether data were crude or *ilr*-transformed. As expected, the clay fraction determined by laser in Set1 was poorly related to the NIR spectra (R^2^ = 0.45-0.64) whether the PSD was expressed as *ilr* or % (Table 3). Combining laser and Bouyoucos methods in Set2 improved the clay predictions with R^2^ values of 0.85-0.91. In Set3, PSD predictions were more accurate with the 2-point 2-h Bouyoucos method (R^2^ = 0.49-0.79) than with the multi-point Bouyoucos method (R^2^ = 0.44-0.70). Modeling Set4 showed that NIR was less accurate (R^2^ = 0.40-0.71) compared to MIR (R^2^ = 0.45-0.80). Accuracy was higher with NIR-4X (R^2^ = 0.87-0.95) than with NIR-2X (R^2^ = 0.69-0.92) (Set5). Models for Set6 showed that data expressed as *ilr* or % (R^2^ = 0.76-0.94) for interpolated PSD were similarly accurate to the slope and intercept from the relationship between settling time and suspension density (R^2^ = 0.84-0.92). Several features in Set7 improved model accuracy (R^2^ = 0.86-0.91) compared to no features at all (R^2^ = 0.82-0.97). The total carbon content contributed the most to the increase in accuracy, followed by colors and oxalate extracts. Finally, the carbon content showed higher performance with raw data (R^2^ = 0.63-0.89) than with *ilr*-transformed data (R^2^ = 0.28-0.84). Compared to a model without features (R^2^ = 0.97), the reconstituted bulk density (R^2^ = 0.99), color (R^2^ = 0.96), oxalate (R^2^ = 0.98) and Mehlich-3 (R^2^ = 0.98) extracts were similar to the accuracy of the NIR model. For the sand fraction, the NIR model calibrated against the sand-sieving method showed higher accuracy (R^2^ = 0.66) compared to the Bouyoucos 2-point 2-h method (R^2^ = 0.53).

**Table 3.**
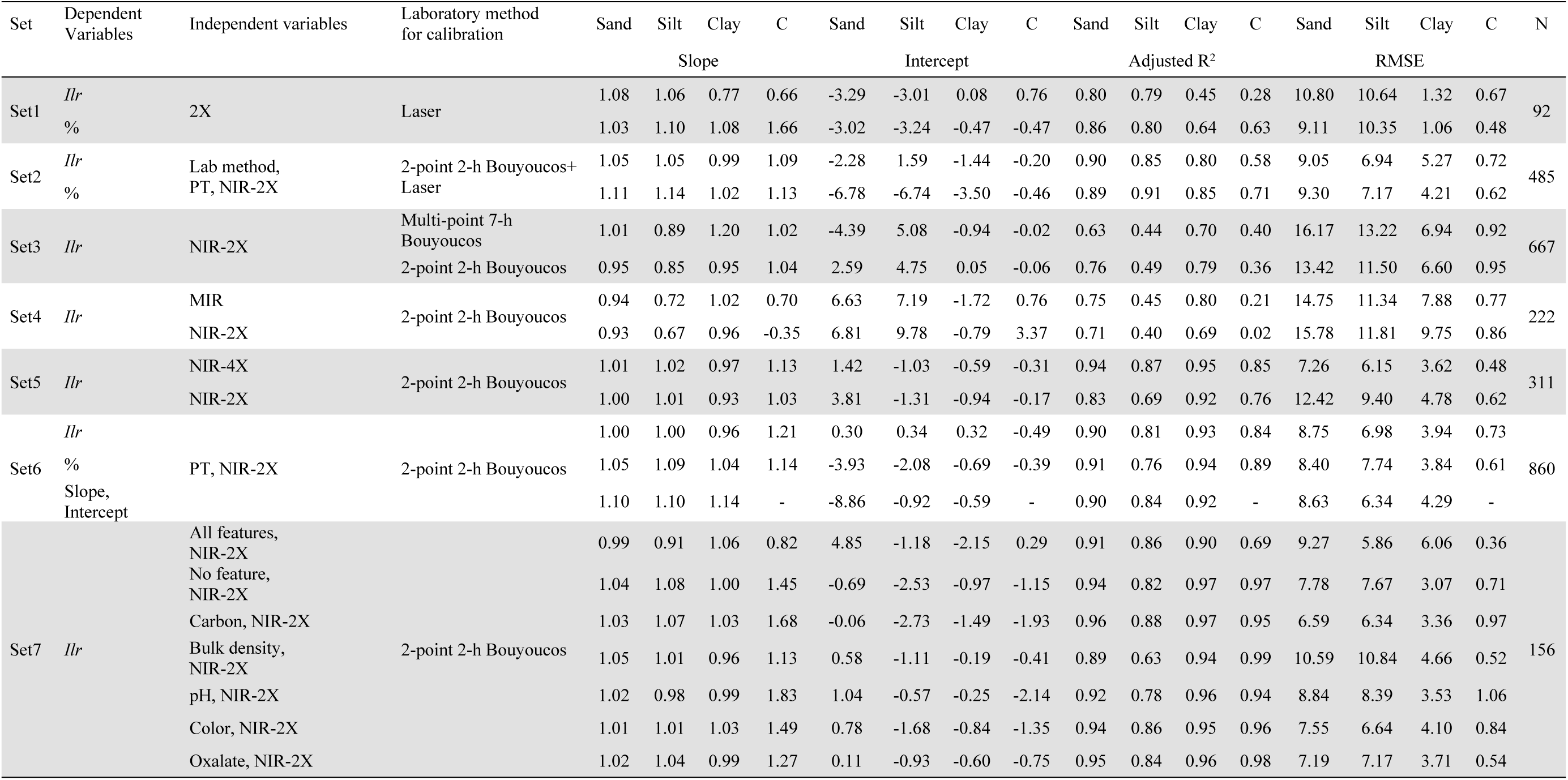
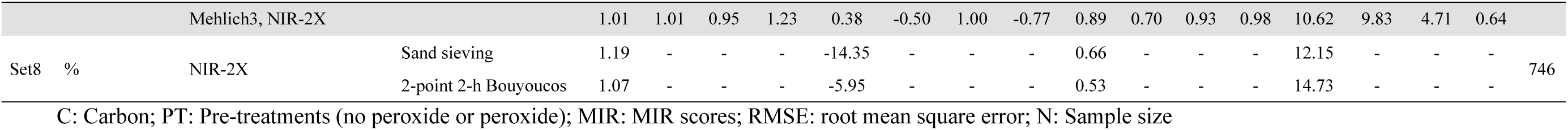
Accuracy, slope, intercept, and number of observations of models

## 4 Discussion

### 4.1 Methodologies

The IR technologies are generally calibrated against a single methodology. In this paper, several methodologies were IR-tested on comparable soil samples. The present paper presented IR-SGB results across several methodologies and features that are rarely addressed altogether in the literature. Machine learning integrated the available information on pre-treatments, methods and features to predict soil texture at low cost in routine laboratories.

#### 4.2.1 Bouyoucos methods

The peroxide pre-treatment tended to increase the clay and silt fractions at the expense of the sand fraction, indicating that clay- and silt-size particles were freed from the clay- or silt-organic matter complexes in micro-aggregates [48]. While metals such as Fe are sequestered by soil organic matter [49] or associated with it [50], iron may be freed from organic matter using hydrogen peroxide to form reactive iron hydroxide [51] that may, conversely, positively impact soil aggregation [52]. Peroxide-treated soils may thus require additional dispersion prior to sedimentation [53]. Nevertheless, the 2-point 2-h Bouyoucos with and without peroxide pre-treatments showed comparable results or minor differences for the [clay | silt,sand] and [silt | sand] balances.

#### 4.2.2 Laser method

The Mie theory considers soil particles to be spherical (Malvern Instruments Ltd., 2007). Parameters such as the refractive index of the medium, the real or imaginary refractive index and the density could be included to improve accuracy. The 2-min sonicated laser method and the sand sieving method returned closed results. The 40 kHz sonication for a duration of 2 min appeared to be suitable for sandy soils [54]. Variants include 36 kHz for 3 min [55] and 30 kHz for 30 min [2]. Although fragile quartz grains may be broken using ultrasonication [56], the [silt | sand] balance for the Bouyoucos method showed results close to the 2-min sonicated laser method.

According to Storti and Balsamo (2010), the high-strength materials are not as affected by the procedures as the low-strength materials. Using water as the dispersing agent and 1750-, 700-rpm for the pump and stirrer speeds, Sochań et al. (2012) obtained R^2^ values of between 0.67 and 0.95 for high-strength materials in silt loamy to sandy soils, values close to our raw data results (R^2^ = 0.64-0.86). For clayed soils, a surfactant or solvent may replace tap water [59]. As small particles increase in number and combine into sand-sized aggregates, pre-treatments may be required and adjusted to soil specificities. There is no standard procedure to disperse soil samples for the laser method because pre-treatment is soil specific.

### 4.3 Spectral data modeling

In this study, we used subsets ≥ 92 samples and GBM as the machine learning method for particle-size prediction compared to suggested subsets ≥ 130 samples for cLHS and FCMS [60]. Here, only Set1 for laser samples did not meet that criterion. The Boosted Regression Trees is another suitable ML method for complex predictions [41] and is relatively fast compared to random forest and neural network algorithms. Model accuracies in validation were in the range reported in the literature where R^2^ values have been found to vary between 0.46 and 0.94 for clay [10,61,62] and between 0.53 and 0.82 for sand. More generally, the R^2^ values may vary widely between 0.05 and 0.84 [63,64].

### 4.4 Prediction accuracy

Because the 2-point 2-h Bouyoucos with and without pre-treatment showed comparable results, they could be combined for IRS calibration with method and treatment information inputs. Hence results of soil surveys could provide a large database for IRS calibration purposes.

Laser methods produced results with lower accuracy compared to the Bouyoucos methods, due to underestimation of the clay fraction. The [silt | sand] balance from the laser method, however, was close to that of the 2-point 2-h Bouyoucos method, despite higher silt and sand percentages for the laser method, indicating advantage for log-ratioing. Higher accuracies from the laser method were obtained by Blott and Pye (2006) across a wide range of soils, sediments and powders. Zobeck (2004) related results from a LS-230 laser diffraction particle size analyzer to those of the pipette method and obtained R^2^ values of 0.97, 0.99, and 0.99 for the < 2-, < 50-, and < 100 –µm in non-calcareous soils, using a shape factor of 0.2 compared to the default 1.0 shape factor for the laser method. Nevertheless, we found that the laser method did not produce results as consistent as the Bouyoucos method for clay predictions by IRS, indicating regional, soil-specific, calibration.

Mehlich-3 extracts, reconstituted bulk density and pH did not improve prediction accuracy. Soil features such as total carbon content, colors and amorphous materials (oxalate extracts) increased model prediction accuracy. Prediction accuracy of carbon content could perform with features as reconstituted bulk density as well as Mehlich-3, colors and oxalate extracts.

Using suspension density function parameters instead of arbitrary settling times did not increase the accuracy of PSD predictions but provided a uniform base to run NIR models as various settling times have direct impact on predictions. Then, in Set3, we observed that 0.67-min and 2-h settling times were more accurate than 0.75-min and 7-h periods for sand and clay NIR predictions.

In the present study, MIR spectra were more accurate than NIR for sand, silt and clay determinations. The NIR method can provide accurate prediction for clay as it has been also found to be accurate for cation exchange capacity (CEC) with R^2^ values of 0.82 [41] and 0.81 [61]. On the same direction, Viscarra Rossel et al. (2006) concluded that MIR was more suitable than NIR for texture and carbon determination, due to higher incidence of spectral bands combined with higher intensity and specificity of the signal compared to NIR. To further support NIR calibration and model accuracy, the R-coded GBM machine learning model used in the present study came across several soil textural classes, carbon contents and features that have not been addressed simultaneously in past research.

## 5 Conclusions

In this paper, IRS inaccurately predicted the PSD’s clay fraction using laser methods. However, IRS accurately predicted PSD against sedimentation and sieving methods after adding soil features such as color, total carbon content and concentration of amorphous materials related to soil genesis and classification. Soil pre-treatments and the need for dispersing agents could be adjusted to the nature and concentration of binding agents for silt-, clay-size particles and fine particles adhering to sand particles. Combined with method and treatment as features, post-screening total carbon content and color routinely determined in service laboratories can improve IRS accuracy for mineral soils. Features as reconstituted bulk density and Mehlich-3 extracts could be added as features for higher-C soils.

The GBM returned similar results whether particle-size data were analyzed as raw pre-determined settling times (percentages) or as *ilr*-transformed percentages. The GBM returned similar accuracies using the slope and intercept of the log-log relationship between settling time and suspension density, *ilr*-transformed percentages or raw percentages. The MIR and NIR 4X gain methods performed better than the NIR 2X gain method. However, additional features increased NIR 2X predictability. Modeling the log-log relationship between settling time and suspension density provided greater flexibility in the choice of soil-specific settling times.

## Acknowledgements

This project was funded by the Natural Sciences and Engineering Council of Canada (CG-2254 and CRDPJ 385199-09), Cultures Dolbec Inc., Groupe Gosselin FG, Prochamps Inc., and Ferme Daniel Bolduc Inc. We thank Catherine Tremblay, Gilles Tremblay, Lucie Grenon, Mario Laterrière, Marie-Hélène Lamontagne, Nicolas Samson, Marie-Ève Tremblay, Lotfi Khiari and Daniel Marcotte for providing soil samples, and Jonathan Lafond for R coding of the laser method.

## Supplementary data

The R code elaborated by Elizabeth Parent and Serge-Etienne Parent is available at https://git.io/fjMIK.

